# Cortisol homeostasis and glucocorticoid signaling are altered in *FMR1* full mutation and premutation carriers

**DOI:** 10.64898/2026.10.20.759351

**Authors:** Heena Mushtaq, Mudasir Manzoor Reshi, Syed Karrar Hussain, Adan Hafiz, Abrar Qurashi

## Abstract

Fragile X syndrome (FXS) and fragile X-associated tremor/ataxia syndrome (FXTAS) are caused by CGG-repeat expansions in the *FMR1* gene. Although the genetic mechanisms underlying these disorders are well established, the contribution of glucocorticoid signaling pathways and neuroendocrine regulation remains unclear. We investigated whether *FMR1* CGG-repeat expansions are associated with alterations in cortisol homeostasis and glucocorticoid signaling pathways.

A clinically referred cohort of 891 individuals was screened for *FMR1* CGG-repeat expansions using conventional and methylation-specific polymerase chain reaction (PCR). Molecular screening identified full mutation (FM) alleles in five males and five females and premutation (PM) alleles in seven males and three females. Endocrine, transcriptional, and epigenetic analyses were subsequently performed in 20 molecularly confirmed carriers (10 PM and 10 FM). Hypothalamic-pituitary-adrenal (HPA) -axis activity was assessed by measuring plasma cortisol concentrations using enzyme-linked immunosorbent assay (ELISA), and the expression of stress-responsive genes (*NR3C1* and *FKBP5*) was quantified by real-time PCR. Epigenetic regulation was further evaluated by assessing *NR3C1* promoter methylation using methylation-specific PCR. To investigate the direct effects of expanded CGG repeats on glucocorticoid signaling independent of systemic influences, complementary in vitro studies were performed in HEK293 cells transiently expressing the expanded CGG repeats. Cells were subsequently subjected to a 12-hr dexamethasone challenge, followed by analysis of *NR3C1* and *FKBP5* expression.

In this preliminary cohort of 20 molecularly confirmed carriers, PM and FM carriers exhibited elevated cortisol concentrations compared with controls. Carriers showed reduced *NR3C1* expression, increased *FKBP5* expression, and increased methylation of the *NR3C1* promoter. Correlation analysis revealed significant relationships between cortisol, *NR3C1,* and *FKBP5* expression. Expression of expanded CGG repeats in HEK293 cells reproduced the glucocorticoid signaling signature observed in patient samples, characterized by reduced *NR3C1* and increased *FKBP5* expression. Following dexamethasone challenge, *FKBP5* expression increased in all experimental groups, with the greatest relative induction observed in CGG90-expressing cells, while *NR3C1* expression remained substantially lower in CGG90-expressing cells than in wild-type cells. These findings indicate altered glucocorticoid-responsive transcription in the CGG-repeat-expressing cellular model. Although these findings require validation in larger independent cohorts, they identify the *NR3C1*-*FKBP5* axis as a potential molecular component of stress-axis dysfunction in fragile X-associated disorders and provide a foundation for future mechanistic studies.

## Introduction

Fragile X syndrome (FXS) is the most common inherited cause of intellectual disability and autism spectrum disorder, resulting from the expansion of a CGG trinucleotide repeats within the 5′ untranslated region of the *FMR1* gene (1,2). Individuals carrying more than 200 CGG repeats typically exhibit FM alleles that become hypermethylated, leading to transcriptional silencing of *FMR1* gene and loss of fragile X messenger ribonucleoprotein (FMRP). In contrast, PM alleles containing 55–200 CGG repeats remain transcriptionally active and are characterized by elevated *FMR1* mRNA levels, which contribute to RNA-mediated toxicity and repeat-associated non-AUG (RAN) translation of potentially toxic polyglycine-containing proteins (3). These distinct molecular mechanisms underlie a spectrum of disorders collectively referred to as *FMR1*-associated disorders, including fragile X syndrome (FXS) and fragile X-associated tremor/ataxia syndrome (FXTAS) (4). Considerable progress has been made in understanding the neurological and molecular consequences of *FMR1* CGG-repeat expansion. FMRP deficiency disrupts synaptic development, neuronal plasticity, and translational regulation, whereas expanded CGG-repeat RNA promotes the sequestration of RNA-binding proteins, cellular stress responses, and neurodegeneration (5,6). However, the biological effects of *FMR1* expansion may extend beyond the nervous system. Individuals with FXS frequently exhibit anxiety, emotional dysregulation, sensory hypersensitivity, and abnormal physiological responses to stress, whereas patients with FXTAS commonly experience anxiety, sleep disturbances, and autonomic dysfunction (7,8).

The hypothalamic–pituitary–adrenal (HPA) axis is the principal neuroendocrine system that regulates physiological and psychological stress responses. Activation of this pathway culminates in the secretion of glucocorticoids, principally cortisol, which regulates metabolism, immune function, neuronal activity, and adaptive stress responses (9). Cortisol signaling is mediated through the glucocorticoid receptor encoded by *NR3C1*, whose activity is tightly controlled by feedback regulators such as *FKBP5*(10,11). Several lines of evidence suggest that *FMR1* expansion may lead to hyperactivation of the HPA axis and impaired glucocorticoid feedback. Elevated cortisol concentrations have been described in PM-carrier mothers experiencing chronic caregiving stress (12). Other investigations, however, have reported normal basal cortisol levels, blunted cortisol awakening responses, or stress-dependent alterations that varied according to CGG repeat size rather than generalized hypercortisolemia (13,14). These apparently conflicting observations collectively indicate that basal cortisol concentrations alone are insufficient to characterize glucocorticoid dysregulation in *FMR1*-associated disorders. Notably, animal studies have implicated altered HPA-axis regulation and abnormal stress responsiveness in *FMR1*-deficient mouse models, including impaired glucocorticoid feedback, altered hypothalamic activation, and behavioral hypersensitivity to stress (15). Direct measurements of circulating cortisol levels, *NR3C1*/*FKBP5* expression, and *NR3C1* promoter methylation in genetically confirmed human PM and FM carriers have not yet been reported. This gap limits the mechanistic understanding of whether glucocorticoid dysregulation is a primary consequence of *FMR1* expansion or a secondary adaptation to chronic stress.

In this study, we investigated whether *FMR1* CGG-repeat expansion is associated with alterations in cortisol homeostasis, glucocorticoid signaling pathways, and epigenetic regulation in genetically confirmed PM and FM carriers. We quantified circulating cortisol concentrations, assessed the expression of the glucocorticoid-signaling genes *NR3C1* and *FKBP5*, examined *NR3C1* promoter methylation, and evaluated the cell-autonomous effects of expanded CGG-repeat expression in an HEK293 cell model. By integrating endocrine and molecular markers in FM, PM carriers, and neurologically normal controls, this study sought to elucidate stress-related mechanisms underlying clinical variability and to explore potential intergenerational modulation of stress biology in fragile X–associated disorders.

## Results

### Cohort ascertainment

We established a clinically referred cohort of individuals with suspected *FMR1*-associated disorders and molecularly characterized the *FMR1* CGG-repeat expansion status. The study cohort comprised 891 participants, including 541 males and 350 females, from independent, non-consanguineous families residing in the central, northern, and southern regions of Kashmir (Fig. 1A). After referral, all participants underwent detailed clinical assessments and were stratified into phenotypically affected and unaffected groups. Among the males, 232 were phenotypically affected, and 309 were phenotypically unaffected. Among the females, 135 were classified as affected and 215 as unaffected (Fig 1B). The clinical manifestations of the affected individuals were consistent with the established FXS phenotype.

**Figure 1.**
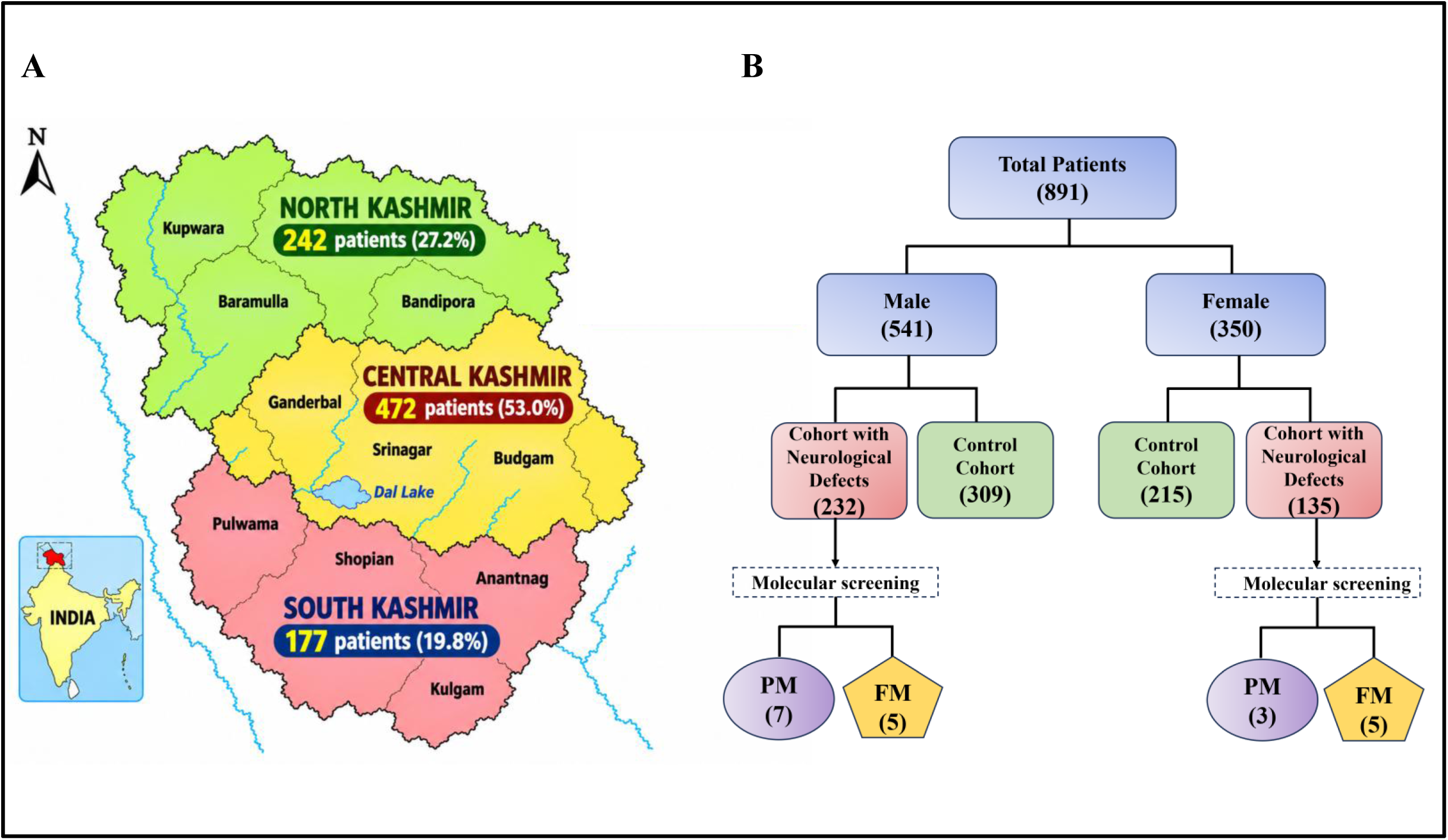
Geographic distribution and sex-wise classification of study participants recruited for *FMR1* PM and FM screening. (A) Map showing the geographical distribution of 891 study participants across the Kashmir region; North Kashmir (242; 27.2%), Central Kashmir (472; 53.0%), and South Kashmir (177; 19.8%). (B) Flow diagram illustrating the sex-wise distribution of participants, comprising 541 males and 350 females. Following molecular screening, 7 males and 3 females were identified with an FMR1 premutation (PM), while 5 males and 5 females were identified with an FMR1 full mutation (FM).

### Molecular confirmation of *FMR1* expansions

Molecular characterization of the *FMR1* CGG-repeat status was subsequently performed using conventional PCR (Fig 2A,2B) and methylation-sensitive PCR (MS-PCR) (Fig 2C,2D). FM and PM alleles were identified exclusively in phenotypically affected individuals within this clinically ascertained cohort. The absence of expanded alleles among phenotypically unaffected participants reflects a phenotype-based referral design rather than a biological finding of mutation penetrance. Among males, FM alleles were detected in five affected individuals (0.92%), and PM alleles were identified in seven affected individuals (1.29%). Among females, FM alleles were identified in five affected individuals (1.42%), and PM alleles were identified in three affected individuals (0.85%). Overall, ten FM carriers and ten PM carriers were identified and constituted the primary cohort for subsequent endocrine, transcriptional, epigenetic, and cellular analyses.

**Figure 2.**
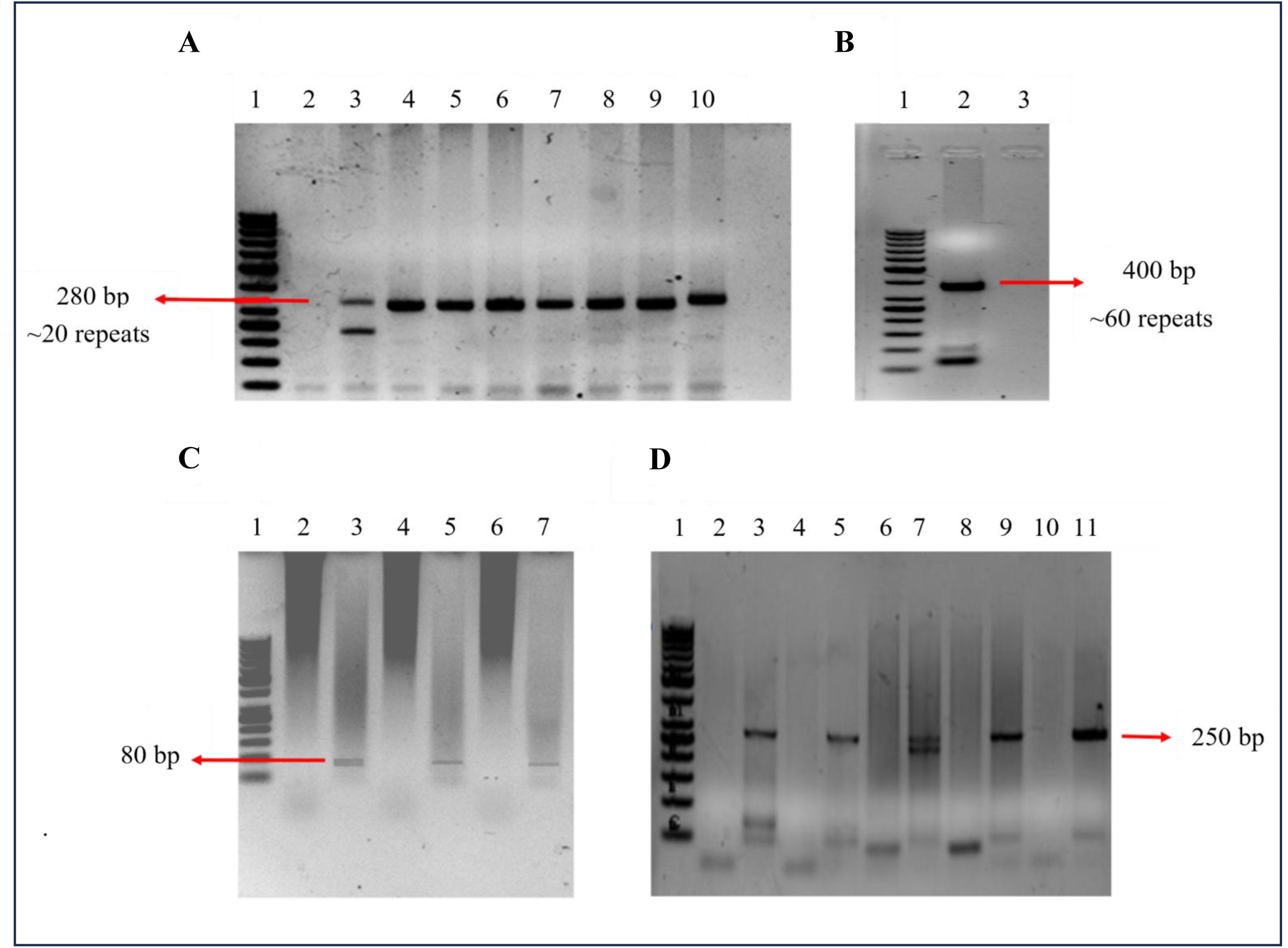
Molecular characterization of *FMR1* CGG-repeat expansions using conventional PCR and methylation-specific PCR (MS-PCR). (A) Agarose gel electrophoresis of conventional PCR amplification of the *FMR1* CGG-repeat region. Lane 1 represents 50 bp DNA ladder; Lane 2 represents suspected male FM subject showing failure of amplification; Lanes 3–10: represents normal individuals. (B) Conventional PCR showing a PM allele. Lane 1 represents 50 bp DNA ladder; Lane 2 represents male PM carrier with an expanded amplicon (∼400 bp); Lane 3 represents NTC. (C) Methylation-specific PCR of male FM subjects following bisulfite modification. Lane 1 represents 50 bp DNA ladder; Lanes 2, 4, and 6 represents amplification with unmethylated-specific primers; Lanes 3, 5, and 7 represents amplification with methylated-specific primers. (D) Methylation-specific PCR of representative normal subjects. Lane 1 represents 50 bp DNA ladder; Lanes 2, 4, 6, 8, and 10 represents methylated-specific primers; Lanes 3, 5, 7, 9, and 11: unmethylated-specific primers.

### *FMR1* expansions are associated with elevated cortisol concentrations

Plasma cortisol, the terminal effector hormone of the HPA axis, was measured to evaluate HPA-axis regulation in PM and FM individuals. Plasma was separated from whole blood by density-gradient centrifugation, and cortisol concentrations were quantified using competitive ELISA. The assay demonstrated adequate sensitivity and reproducibility for precise interpolation of sample values, as confirmed by the standard curve generated by plotting known cortisol concentrations against optical density (OD), which showed the expected inverse relationship between OD and cortisol concentration. As plasma cortisol is a principal effector hormone of the HPA axis and a widely accepted biomarker of systemic glucocorticoid signaling, its concentrations were quantified in controls and molecularly characterized individuals carrying PM and FM alleles. Analysis of log-transformed cortisol concentrations revealed a significant effect of genotype on circulating cortisol levels (one-way ANOVA, P < 0.0001, η² = 0.675). Tukey’s multiple-comparison test demonstrated that both FM and PM carriers exhibited significantly higher cortisol concentrations than control subjects (Control vs. FM: mean difference = −4.609, P < 0.001; Control vs. PM: mean difference = −4.151, P < 0.001), whereas no significant difference was observed between the PM and FM groups (mean difference = 0.4573, P = 0.7840) (Fig. 3A). These findings indicate that dysregulation of the HPA axis is already evident in PM carriers and persists following expansion to the FM, suggesting that elevated glucocorticoid activity is an early and sustained feature of the *FMR1* mutation spectrum.

**Figure 3.**
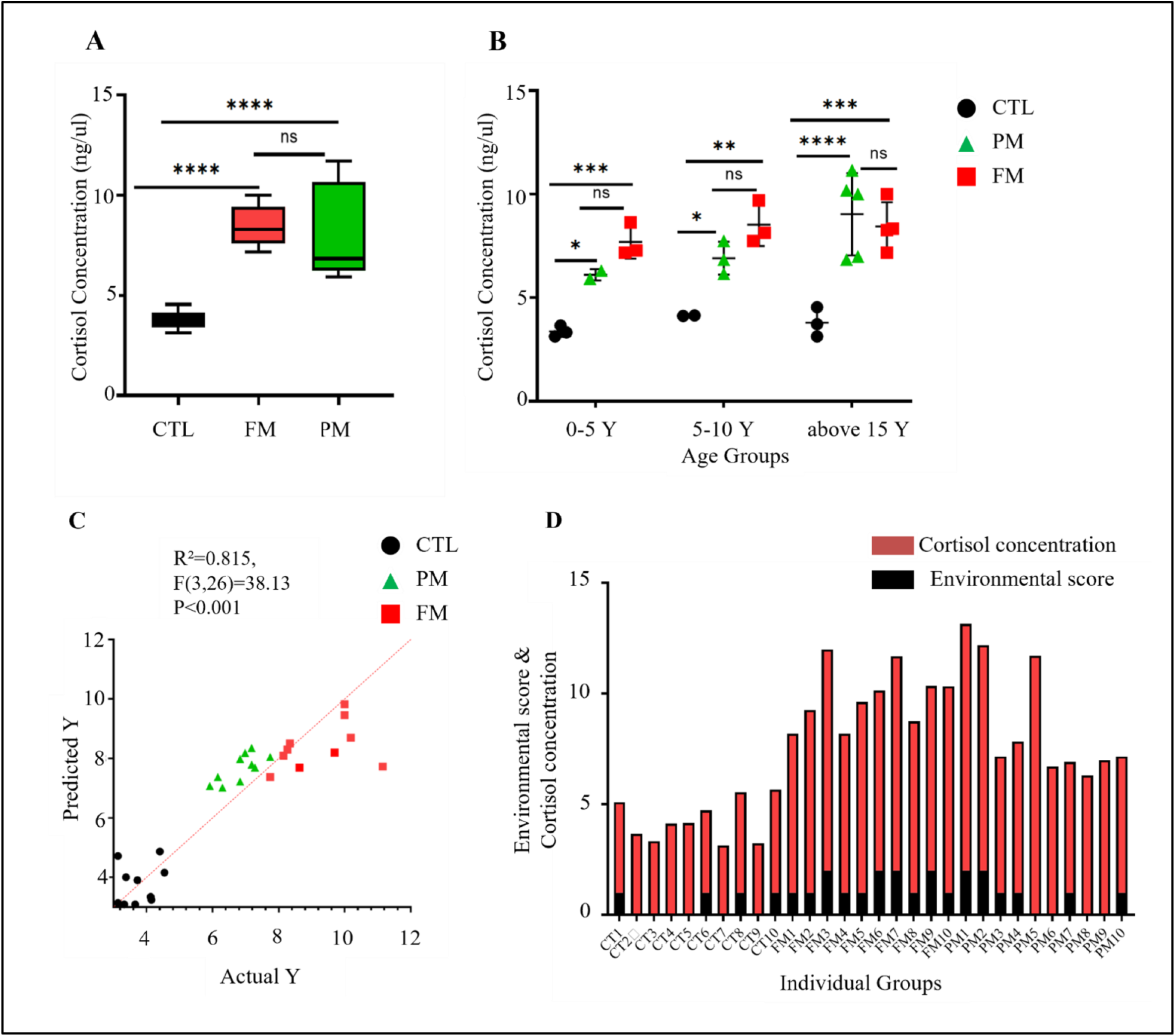
Plasma cortisol concentrations in CTL, PM, and FM individuals. (A) Box-and-whisker plot showing plasma cortisol concentrations in CTL, PM, and FM groups. (B) Age-stratified analysis of plasma cortisol concentrations in CTL, PM, and FM individuals grouped into 0–5 years, 5–10 years, and >15 years. (C) Multiple linear regression analysis showing the relationship between observed (Actual Y) and predicted plasma cortisol concentrations after adjustment for age. (D) Individual plasma cortisol concentrations and corresponding environmental stress scores in CTL, PM, and FM participants.

Because age-dependent penetrance is an intrinsic feature of *FMR1***-**associated disorders, participants were further stratified into three age groups (0–5 years, 5–10 years, and >15 years) to determine whether genotype-associated differences in cortisol concentrations persisted across different stages of development (Fig. 3B). In the 0–5-year age group, both PM and FM individuals exhibited significantly higher cortisol concentrations than controls (Control vs. PM: mean difference = −2.736, P = 0.0465; Control vs. FM: mean difference = −4.328, P = 0.0006), whereas no significant difference was observed between PM and FM groups (P = 0.3134). A similar pattern was observed in the 5–10-year age group, where cortisol concentrations remained significantly elevated in both PM (mean difference = −2.778, P = 0.0429) and FM individuals (mean difference = −4.392, P = 0.0016) compared with controls, with no significant difference between PM and FM groups (P = 0.2310). In participants older than 15 years, cortisol concentrations were markedly elevated in both mutation groups relative to controls (Control vs. PM: mean difference = −5.231, P < 0.0001; Control vs. FM: mean difference = −4.647, P = 0.0001), while PM and FM groups again did not differ significantly (P = 0.7380). Notably, PM individuals in this age group exhibited the highest mean cortisol concentration among all study groups, suggesting that HPA-axis dysregulation becomes increasingly pronounced with advancing age.

To determine whether these genotype-associated differences were independent of age, multiple linear regression analysis was performed using plasma cortisol concentration as the dependent variable, genotype as the independent variable, and age as a continuous covariate (Fig. 3C). The overall regression model was highly significant (F (3, 26) = 38.13, P < 0.0001) and explained 81.5% of the variance in plasma cortisol concentrations (R² = 0.815). After adjustment for age, both PM (β = 4.029, 95% CI: 3.027–5.030, P < 0.0001) and FM carriers (β = 4.647, 95% CI: 3.641–5.654, P < 0.0001) continued to exhibit significantly higher plasma cortisol concentrations than controls. Age also emerged as an independent positive predictor of cortisol concentration (β = 0.0508, 95% CI: 0.0194–0.0821, P = 0.0026), indicating that circulating cortisol concentrations increased modestly with advancing age irrespective of genotype. Collectively, these findings demonstrate that elevated cortisol concentrations in *FMR1* mutation carriers are not solely attributable to age-related physiological variation but represent genotype-associated dysregulation of the HPA axis that becomes more pronounced with increasing age.

Environmental stress scores remained low and comparable across the control, PM, and FM groups and showed no significant association with genotype (Fig. 3D). Similarly, environmental stress scores were not associated with circulating cortisol concentrations. Within the scope of the variables evaluated, these findings suggest that the observed association between *FMR1* expansion status and altered cortisol concentrations was not explained by the environmental stress measures assessed using the study questionnaire.

### Altered glucocorticoid signaling gene expression accompanies cortisol dysregulation

To investigate the molecular pathways associated with elevated cortisol concentrations, the expression of key glucocorticoid-signaling genes was assessed in peripheral blood. We focused on *NR3C1*, which encodes the glucocorticoid receptor responsible for glucocorticoid-mediated negative feedback regulation, and *FKBP5*, a well-established negative regulator of glucocorticoid receptor signaling.

Age-stratified analysis demonstrated a consistent reduction in NR3C1 expression in both PM and FM carriers compared with age-matched controls (Fig. 4A). In the 0–5-year age group, *NR3C1* expression was reduced to 0.51-fold in PM carriers (*P* = 0.0213) and 0.57-fold in FM carriers (*P* = 0.0237) relative to controls. Similar reductions were observed in the 5–10-year age group, with expression levels of 0.56-fold in PM (*P* = 0.0196) and 0.44-fold in FM (*P* = 0.0030). In individuals older than 15 years, *NR3C1* expression remained significantly decreased, reaching 0.65-fold in PM (*P* = 0.0492) and 0.51-fold in FM carriers (*P* = 0.0050). No significant differences in *NR3C1* expression were observed between the PM and FM groups across any age category (*P* > 0.05), indicating that reduced glucocorticoid receptor expression is established at the premutation stage and persists following expansion to the full mutation.

**Figure 4.**
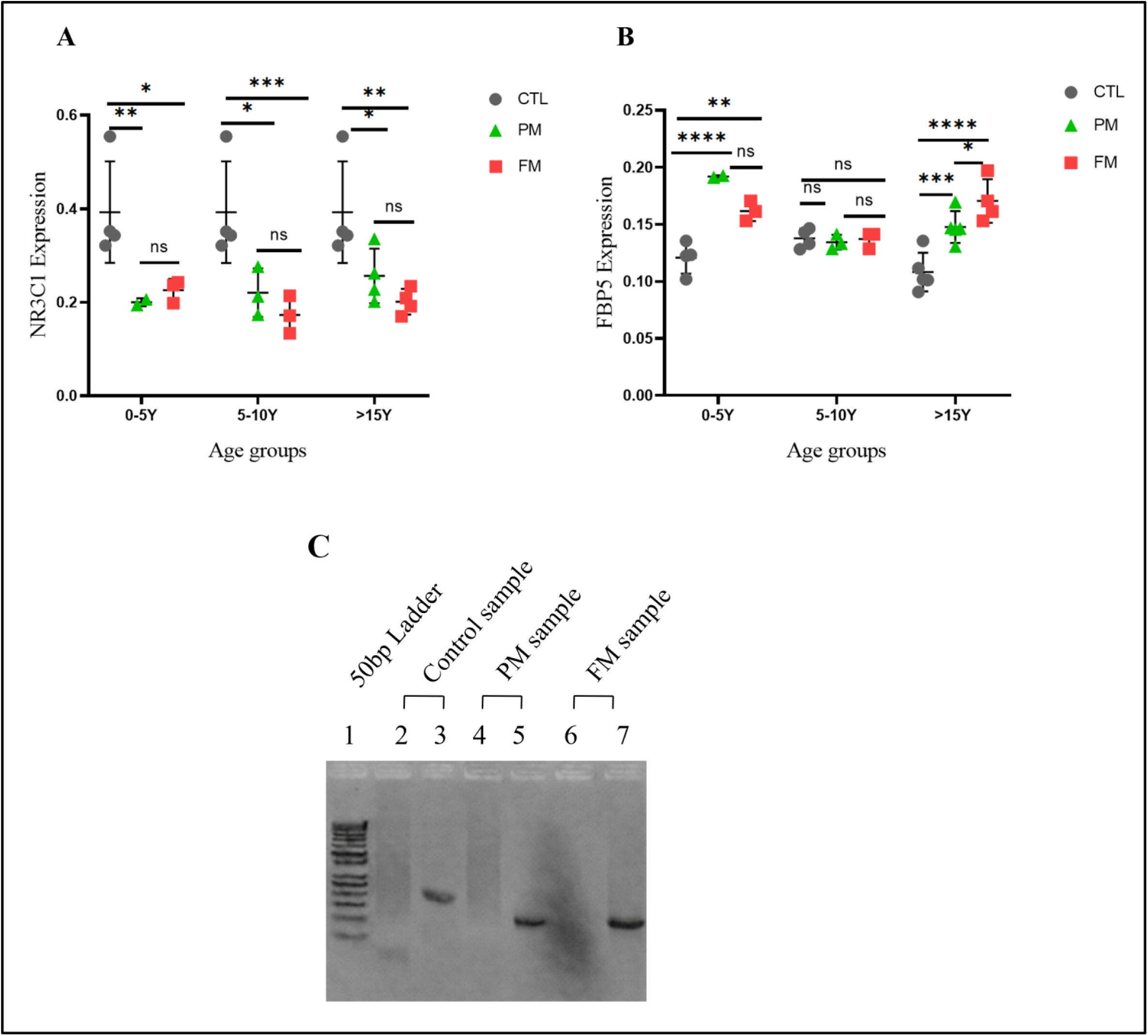
Age-wise expression of *NR3C1* and *FKBP5* genes and methylation-specific PCR analysis of the *NR3C1* promoter in CTL, PM), and FM individuals. (A) Scatter plot showing age-stratified relative *NR3C1* gene expression in CTL, PM, and FM groups categorized into 0–5 years, 5–10 years, and >15 years. (B) Scatter plot showing age-stratified relative *FKBP5* gene expression in CTL, PM, and FM groups categorized into 0–5 years, 5– 10 years, and >15 years. Data are mean ± SEM from three biological replicates. (C) Agarose gel electrophoresis of methylation-specific PCR (MSP) products showing the methylation status of the *NR3C1* promoter. Lane 1 represents 50 bp DNA ladder; Lanes 2 and 3 represent control sample amplified with the unmethylated primer set(214bp); Lanes 4and 5 represent PM samples amplified with the methylated primer set (150bp); Lanes 6 and 7 represent FM samples amplified with the methylated primer set(150bp).

In contrast, *FKBP5* expression exhibited an overall increase in mutation carriers, although the magnitude of induction varied across age groups (Fig. 4B). In the 0–5-year age group, *FKBP5* expression increased to 1.59-fold in PM carriers (*P* < 0.0001) and 1.34-fold in FM carriers (*P* = 0.0013), with PM carriers showing significantly higher expression than FM carriers (*P* = 0.0486). In in individuals older than 15 years, *FKBP5* expression was again significantly elevated, reaching 1.37-fold in PM carriers (*P* = 0.0002) and 1.58-fold in FM carriers (*P* < 0.0001), with FM carriers exhibiting significantly higher expression than PM carriers (*P* = 0.0422).

The combined pattern of reduced *NR3C1* expression and increased *FKBP5* expression is consistent with impaired glucocorticoid receptor signaling and dysregulation of the glucocorticoid feedback pathway. Although these alterations were observed consistently across age groups, the relatively small sample size (n = 20 carriers) and exploratory nature of the study warrant validation in larger, independent cohorts. Furthermore, functional studies are required to determine whether these transcriptional changes directly translate into altered glucocorticoid responsiveness.

### *NR3C1* promoter methylation is associated with altered receptor expression

Given the observed reduction in *NR3C1* transcript levels and elevation of *FKBP5* expression, we investigated whether epigenetic alterations contributed to the altered glucocorticoid signaling in carriers of *FMR1* expansion. Previous studies on chronic stress-related disorders have linked DNA methylation changes at glucocorticoid signaling loci to altered stress responses and endocrine dysfunction.

In our primary cohort, both PM and FM carriers exhibited greater *NR3C1* promoter methylation than controls (Fig. 4C), a finding that requires replication using adequately powered, independent samples. Increased methylation was correlated with reduced *NR3C1* transcript abundance, elevated cortisol concentrations, and increased CGG-repeat length, although a causal relationship was not established, consistent with the potential contribution of epigenetic alterations to altered glucocorticoid signaling. Functional studies are required to determine whether methylation changes actively mediate abnormalities in glucocorticoid signaling.

### Integrated correlation analyses revealed coordinated alterations in the glucocorticoid signaling pathway

To investigate the relationship between HPA-axis activity and stress-responsive gene expression, Spearman’s rank correlation analysis was performed between plasma cortisol concentrations and the expression of *NR3C1* and *FKBP5*. Cortisol concentration showed a significant negative correlation with *NR3C1* expression (r = −0.6916, 95% CI: −0.8543 to −0.4056, *P* < 0.0001), indicating that individuals with elevated cortisol levels exhibited reduced glucocorticoid receptor expression (Fig. 5A). This finding is consistent with impaired glucocorticoid receptor-mediated negative feedback on the HPA axis. A significant inverse correlation was also observed between *NR3C1* and *FKBP5* expression (r = −0.7900, 95% CI: −0.9294 to −0.4533, *P* = 0.0007), demonstrating that lower *NR3C1* expression was associated with higher *FKBP5* expression (Fig. 5B). Conversely, plasma cortisol concentration was positively correlated with *FKBP5* expression (r = 0.6974, 95% CI: 0.3684–0.8712, *P* = 0.0004), indicating that increased systemic glucocorticoid activity was accompanied by enhanced expression of this negative regulator of glucocorticoid receptor signaling (Fig. 5C). Collectively, these findings demonstrate coordinated dysregulation of the *NR3C1*–*FKBP5* signaling axis in *FMR1* mutation carriers.

**Figure 5.**
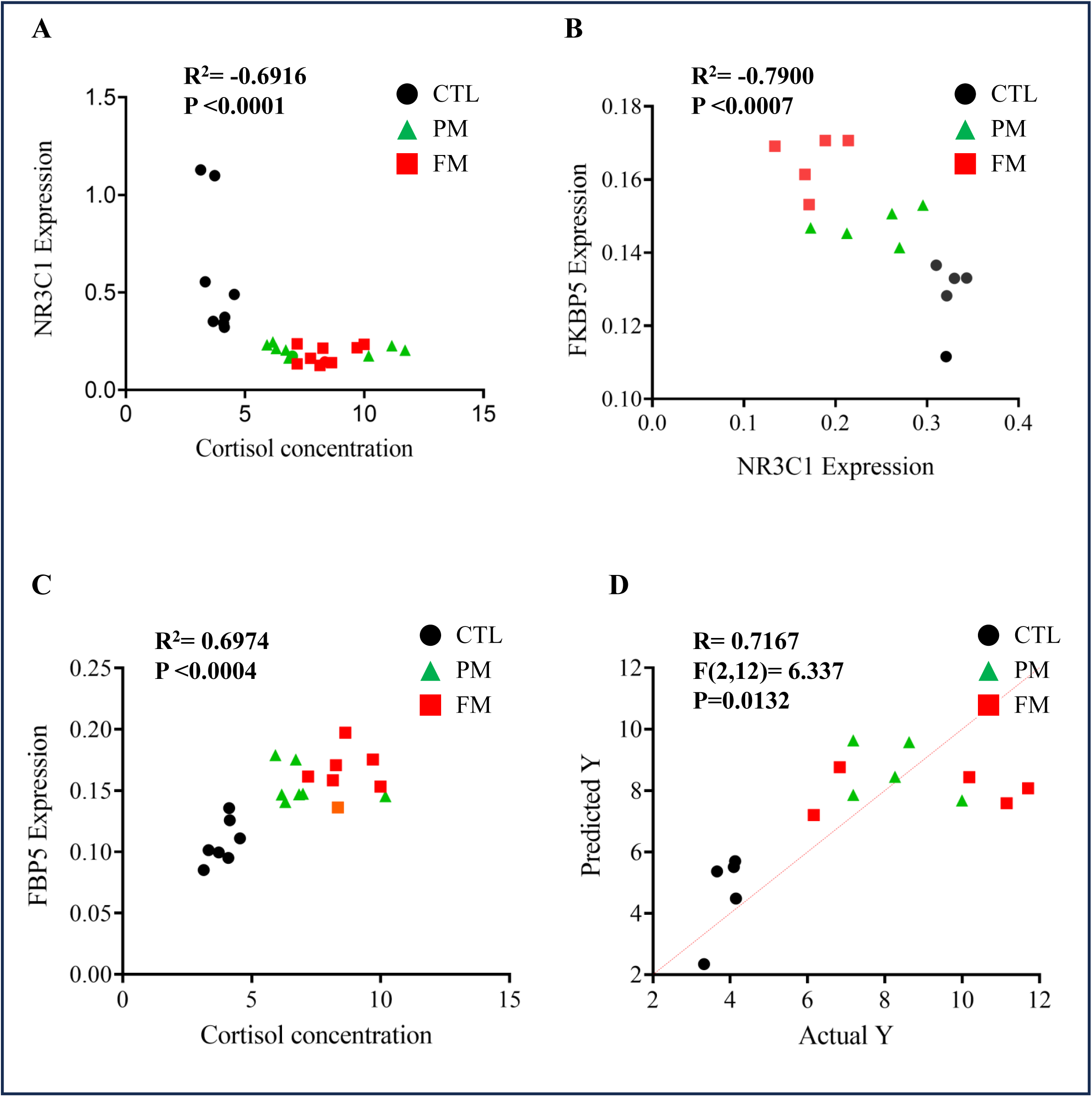
Correlation and multiple regression analyses of cortisol concentration and glucocorticoid signaling markers in CTL, PM, and FM. (A) Scatter plot showing the negative correlation between cortisol concentration and *NR3C1* expression. (B) Scatter plot showing the inverse relationship between *NR3C1* and *FKBP5* expression. (C) Scatter plot demonstrating the positive correlation between cortisol concentration and *FKBP5* expression. (D) Predicted versus observed cortisol concentrations generated from the multiple linear regression model using *NR3C1* and *FKBP5* expression as predictor variables. The dashed red line represents the line of identity (predicted = observed). Correlation analyses were performed using Spearman’s rank correlation, and the regression model was generated using multiple linear regression analysis.

To further evaluate the independent contribution of stress-responsive genes to circulating cortisol concentrations, multiple linear regression analysis was performed using cortisol concentration as the dependent variable and *NR3C1* and *FKBP5* expression as predictor variables (Fig. 5D). *NR3C1* exhibited a negative regression coefficient (β = −15.33, 95% CI: −31.01 to 0.35) and showed a trend toward significance (*P* = 0.0545), suggesting that reduced glucocorticoid receptor expression may contribute to elevated cortisol concentrations. In contrast, *FKBP5* was not independently associated with cortisol levels after adjustment for *NR3C1* expression (β = 25.83, 95% CI: −76.73 to 128.4, *P* = 0.5933). Although neither predictor reached statistical significance in the multivariable model, the regression analysis, together with the strong pairwise correlations, supports the existence of a coordinated glucocorticoid-signaling network that is disrupted in *FMR1* mutation carriers.

### Expanded CGG repeats reproduce the glucocorticoid signaling dysregulation in HEK293 cells

Although patient-derived analysis has identified elevated cortisol levels and altered glucocorticoid signaling gene expression, these observations could arise either as primary consequences of *FMR1* expansion or as secondary adaptations to chronic endocrine dysregulation. Therefore, we sought to determine whether CGG-repeat expression alone was sufficient to reproduce the observed molecular signature in a cell-autonomous manner. HEK293 cells were selected as a reductionist heterologous system to determine whether the expression of expanded CGG repeats was sufficient to alter glucocorticoid signaling pathways independent of systemic endocrine influences.

HEK293 cells were transfected with constructs containing either wild-type or expanded CGG90 repeat sequences, with an empty vector serving as the transfection control. In the absence of dexamethasone, CGG90-expressing cells exhibited marked downregulation of *NR3C1* expression and increased *FKBP5* expression relative to wild-type cells. Relative expression analysis showed that *NR3C1* expression was approximately 0.18-fold in CGG90-expressing cells compared with wild-type cells, whereas *FKBP5* expression was approximately 1.56**-**fold. These findings indicate that expanded CGG-repeat expression is associated with coordinated alterations in glucocorticoid receptor signaling components, characterized by reduced *NR3C1* expression and increased *FKBP5* expression.

To further examine the glucocorticoid responsiveness of this pathway, cells were subjected to a 12-h dexamethasone challenge following 36 h of transfection. Dexamethasone treatment increased *FKBP5* expression in all three experimental backgrounds. *FKBP5* expression increased from approximately 1.00-fold in untreated wild-type cells to 1.72-fold following dexamethasone treatment. In vector-transfected cells, *FKBP5* expression increased from 1.29-fold to 1.56-fold, whereas CGG90-expressing cells showed the largest response, increasing from 1.56-fold to 2.71-fold relative to untreated wild-type cells. Thus, CGG90-expressing cells demonstrated a pronounced *FKBP5* induction following dexamethasone exposure (Fig 6A).

**Figure 6.**
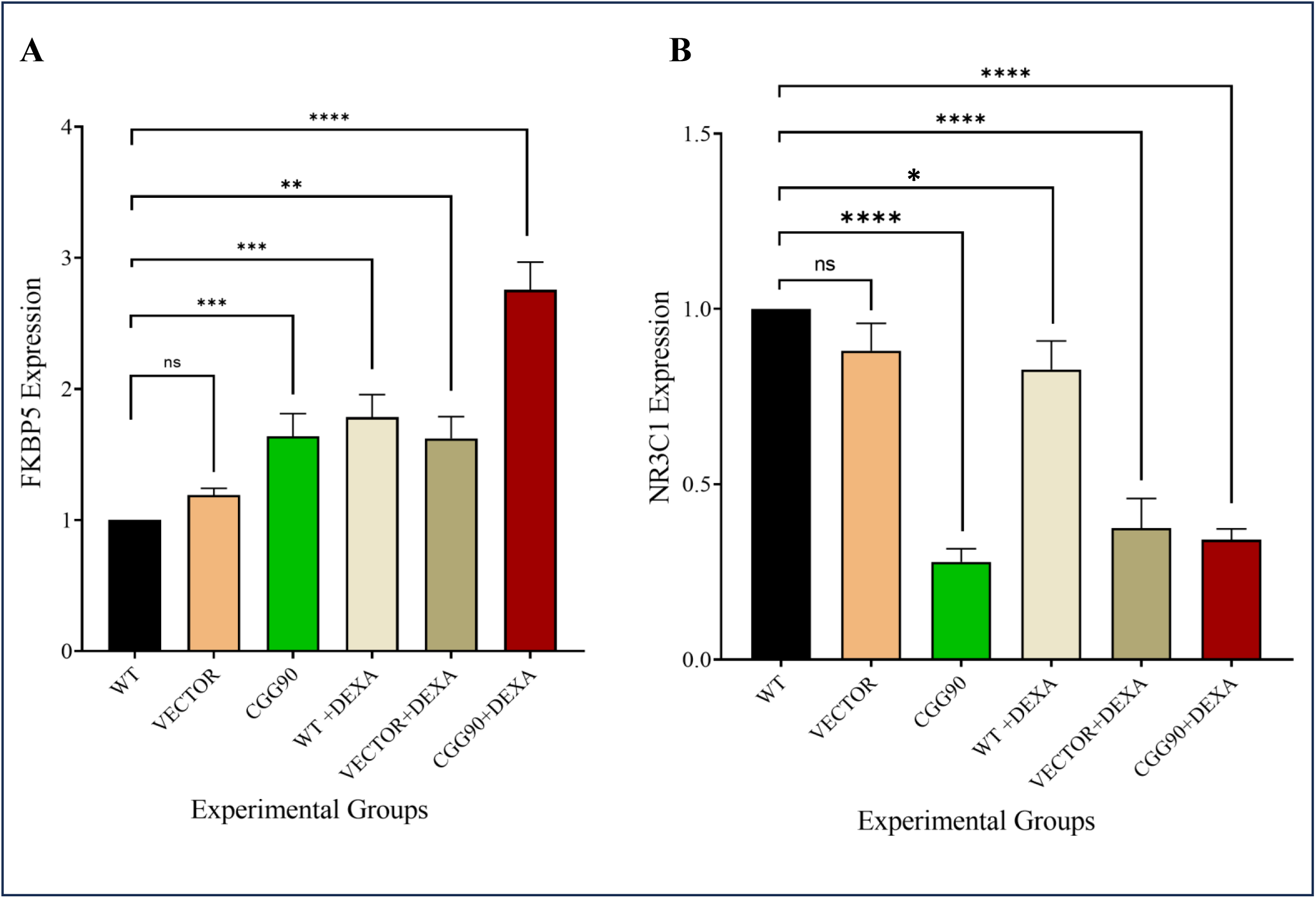
Expanded CGG repeats alter glucocorticoid-responsive gene expression following dexamethasone treatment in HEK293 cells. Relative mRNA expression of (A) *FKBP5* and (B) *NR3C1* was measured by qRT-PCR in WT, vector, and CGG90 cells with or without dexamethasone (Dex). Expression was normalized to actin using the 2⁻ΔΔCt method. Data are mean ± SEM from three biological replicates.

In contrast, *NR3C1* expression decreased following dexamethasone treatment in wild-type and vector-transfected cells, from approximately 1.00-fold to 0.65-fold and from 0.58-fold to 0.26-fold, respectively. In CGG90-expressing cells, *NR3C1* expression increased modestly following dexamethasone exposure, from 0.18-fold to 0.25-fold; however, expression remained substantially below the untreated wild-type level. Thus, dexamethasone exposure did not restore *NR3C1* expression to control levels in CGG90-expressing cells, despite the modest increase following treatment (Fig 6B).

Taken together, these findings demonstrate that expanded CGG-repeat expression is associated with a distinct glucocorticoid signaling phenotype in HEK293 cells, consisting of low basal *NR3C1* expression and elevated *FKBP5* expression, together with a strong *FKBP5* response to exogenous dexamethasone and persistently low *NR3C1* expression. The dexamethasone-response experiment therefore provides additional evidence that CGG-repeat expression can alter glucocorticoid-responsive transcription in a cell-autonomous setting. Because the CGG90 construct models an expanded, transcriptionally active premutation-associated repeat, these findings support the possibility that expanded CGG-repeat RNA contributes directly to dysregulation of glucocorticoid signaling. However, because HEK293 cells represent a non-neuronal heterologous system lacking the physiological HPA-axis context, these observations should be considered proof-of-concept evidence rather than direct evidence of neuronal or systemic pathophysiology.

## Discussion

While isolated alterations in stress reactivity have previously been reported in individuals with *FMR1* expansions, this study is the first to demonstrate that *FMR1* CGG-repeat expansions are associated with coordinated alterations across the endocrine, transcriptional, and epigenetic components of glucocorticoid signaling. Our studies on patient-derived samples and in vitro experiments support the hypothesis that dysregulation of glucocorticoid signaling represents an underrecognized molecular consequence of *FMR1* expansion. The relationship between *FMR1* CGG-repeat expansions and HPA-axis dysfunction remains unclear because previous studies have reported inconsistent alterations in cortisol homeostasis. Elevated cortisol concentrations have been described in PM carrier mothers experiencing chronic caregiving stress, whereas other investigations have reported normal basal cortisol levels, blunted cortisol awakening responses, or stress-dependent alterations that varied according to CGG repeat size rather than generalized hypercortisolemia. These apparently conflicting observations collectively indicate that basal cortisol concentrations alone are insufficient to characterize glucocorticoid dysregulation in *FMR1*-associated disorders.

Our findings help reconcile these discrepancies by demonstrating that the assessment of circulating cortisol alone provides only a partial view of glucocorticoid dysfunction. We showed that elevated plasma cortisol in mutation carriers was accompanied by reduced *NR3C1* expression, increased *FKBP5* expression, and hypermethylation of the *NR3C1* promoter, indicating coordinated impairment of glucocorticoid receptor signaling rather than isolated endocrine dysregulation. Our observations, therefore, suggest that disruption of downstream glucocorticoid receptor regulation may represent a more conserved molecular consequence of expanded CGG repeats.

Our cellular experiments provide complementary mechanistic evidence to support these clinical observations. Expression of expanded CGG90 repeats in HEK293 cells reproduced the molecular signature observed in patient samples, including suppression of *NR3C1* and induction of *FKBP5*, suggesting that expanded CGG-repeat RNA is sufficient to perturb key components of glucocorticoid signaling, independent of systemic endocrine influences. Although glucocorticoid receptor activity was not directly measured, these findings indicate that toxic CGG-repeat RNA intrinsically disrupts the regulators of glucocorticoid responsiveness. Because the heterologous CGG90 construct models an unmethylated, transcriptionally active premutation allele, these experiments specifically isolated the cell-autonomous effects of expanded CGG-repeat RNA and provided proof-of-concept evidence that glucocorticoid pathway dysregulation can arise independently of the physiological HPA axis.

Recent experimental studies have provided additional mechanistic support for this interpretation. Sirois et al. demonstrated that CGG repeats in the *FMR1* gene influence glucocorticoid receptor signaling, receptor localization, and cellular stress responses in developing neurons, suggesting a potential link between *FMR1*-associated molecular changes and altered neuronal responses to glucocorticoids. Similarly, Bhattacharyya et al. showed that expanded CGG-repeat RNA disrupts intracellular RNA localization and stress-responsive pathways during neuronal development. Although these investigations were performed in neuronal systems rather than HEK293 cells, they support the emerging concept that expanded CGG-repeat RNA intrinsically impairs glucocorticoid signaling and are consistent with our observations of reduced *NR3C1* and increased *FKBP5* expression.

The similar neuroendocrine alterations observed in both PM and FM carriers suggest that impaired glucocorticoid signaling may represent a convergent downstream consequence of *FMR1* expansion, despite the distinct pathogenic mechanisms underlying these two mutation classes. Future studies that independently manipulate CGG expression and FMRP abundance within disease-relevant neuronal models are required to distinguish the relative contributions of these pathogenic pathways.

Cross-sectional analyses further demonstrated that cortisol dysregulation was associated with age. Regression analysis incorporating age as a covariate showed that the association between the *FMR1* genotype and circulating cortisol remained significant after adjustment, indicating that the endocrine abnormalities observed in mutation carriers cannot be explained solely by physiological aging. As age-dependent penetrance is an intrinsic feature of *FMR1*-associated disorders, we additionally performed age-stratified analyses across younger (<10 years, 10-15 and above 15 years participants. These complementary analyses suggest that alterations in cortisol homeostasis are associated with *FMR1* expansion independently of age, while also indicating that the magnitude of endocrine dysregulation may increase across clinically relevant stages of disease progression. One possible explanation is that chronic impairment of glucocorticoid feedback progressively destabilizes endocrine homeostasis. Alternatively, age-dependent consequences of CGG-repeat expansion, including RNA toxicity, altered stress granule dynamics, mitochondrial dysfunction, and neuroinflammation, may progressively exacerbate abnormalities in glucocorticoid signaling. Longitudinal studies are required to determine the temporal evolution of these endocrine alterations and establish whether they parallel disease progression.

An important observation was that elevated cortisol concentrations were not significantly associated with the environmental stress measures evaluated in this study. Environmental stress scores were comparable among controls, PM carriers, and FM carriers. However, although the checklist assessed several potential confounding factors, including caregiver-related stress, sleep disturbances, family conflict, recent acute stress, medication use, chronic endocrine or metabolic disorders, and stress associated with blood collection, it may not have fully captured the intensity, duration, or cumulative effects of these exposures. Furthermore, other factors, such as socioeconomic adversity and unmeasured aspects of chronic psychosocial stress, may have contributed to individual variability in HPA-axis activity.

Therefore, although the endocrine abnormalities observed in this study were not associated with the environmental stress measures assessed, residual confounding cannot be entirely excluded. This interpretation is strengthened by our in vitro findings, in which the expression of expanded CGG repeats reproduced the glucocorticoid-signaling signature independently of systemic endocrine and environmental influences, supporting an intrinsic molecular effect of *FMR1* expansion.

Molecular analyses provide a mechanistic framework for the endocrine abnormalities observed in mutation carriers. *NR3C1* encodes the glucocorticoid receptor, the principal mediator of glucocorticoid signaling, and negative feedback regulation within the HPA axis. Reduced *NR3C1* expression is expected to diminish glucocorticoid sensitivity, impair feedback inhibition, and permit sustained cortisol secretion. Consistent with this model, *NR3C1* expression was significantly reduced in both PM and FM carriers and exhibited an inverse correlation with circulating cortisol concentrations. Furthermore, the degree of *NR3C1* downregulation appeared to increase with age, paralleling the higher cortisol concentrations observed in older participants. Although these observations require validation in larger cohorts, they suggest that progressive impairment of glucocorticoid receptor signaling may contribute to age-dependent endocrine dysfunction in individuals with *FMR1* expansion.

In contrast, *FKBP5* expression was significantly elevated in mutation carriers. *FKBP5* is a well-established negative regulator of glucocorticoid receptor signaling that functions within an intracellular feedback loop to reduce receptor activation, ligand affinity, and nuclear translocation. Increased *FKBP5* expression has been implicated in several disorders characterized by impaired glucocorticoid responsiveness, including major depressive disorder, post-traumatic stress disorder, and chronic stress-related conditions. The strong inverse relationship observed between *FKBP5* and *NR3C1* expression suggests that these pathways may cooperate to attenuate glucocorticoid receptor activity in carriers of *FMR1* expansion. Whether increased *FKBP5* expression is a primary pathogenic driver or a compensatory response secondary to chronic glucocorticoid exposure remains unclear and warrants further investigation.

An additional layer of regulation was provided by epigenetic analyses. Mutation carriers exhibited increased methylation of the *NR3C1* promoter, which was consistent with transcriptional repression of the glucocorticoid receptor gene. Previous studies on stress-related neuropsychiatric disorders have shown that *NR3C1* promoter hypermethylation is associated with reduced receptor expression, impaired glucocorticoid feedback, and persistent HPA axis activation. Our findings extend these observations to *FMR1*-associated disorders and suggest that epigenetic silencing of *NR3C1* may contribute to the reduced receptor expression observed in mutation carriers. Although the present study could not establish whether promoter hypermethylation is a primary consequence of expanded CGG repeats or develops secondary to chronic endocrine dysregulation, the concordance between increased *NR3C1* methylation, reduced transcript abundance, elevated *FKBP5* expression, and increased circulating cortisol supports a model of coordinated disruption of glucocorticoid receptor regulation.

Taken together, endocrine, transcriptional, epigenetic, and cellular findings support a unified model in which *FMR1* CGG-repeat expansion perturbs glucocorticoid homeostasis through multiple interacting mechanisms. Expanded CGG-repeat RNA appears to be capable of intrinsically altering glucocorticoid pathway regulators, while reduced *NR3C1* expression, increased *FKBP5* expression, and *NR3C1* promoter hypermethylation collectively diminish glucocorticoid receptor responsiveness and weaken the negative feedback regulation of the HPA axis. Rather than representing isolated molecular abnormalities, these alterations likely reflect interconnected components of a common pathogenic pathway that contributes to endocrine dysregulation in both pre-mutation and full-mutation carriers.

### Limitations

Participants were recruited through phenotype-based clinical referral pathways targeting individuals with intellectual disabilities, autism, or FXTAS-like symptoms. This ascertainment strategy may select carriers with more severe phenotypes, potentially overestimating the prevalence and magnitude of glucocorticoid-signaling abnormalities relative to the broader population of PM/FM carriers, many of whom may be phenotypically unaffected or mildly affected. Population-based screening studies are needed to determine whether cortisol dysregulation is present across the full spectrum of mutation carriers or is enriched in those with overt clinical manifestations. Although 891 individuals were screened, only 20 molecularly confirmed PM/FM carriers were available for endocrine and molecular analyses, which substantially limited the statistical power. Consequently, all molecular findings should be considered exploratory and hypothesis generating, pending replication in larger independent cohorts.

The cross-sectional design precluded causal inferences. While prior studies have reported moderate correlations between the peripheral and brain tissue expression of *NR3C1* and *FKBP5* in some contexts (16) to which blood-based measures capture neuronal glucocorticoid signaling in *FMR1* carriers remains unknown. “Although an environmental and procedural stress checklist was used to assess potential confounding factors, including caregiver-related stress, sleep disturbances, family conflict, recent acute stress, medication use, chronic endocrine or metabolic disorders, and stress associated with blood collection, the assessment may not have fully captured the intensity, duration, or cumulative effects of these exposures. Therefore, residual confounding due to persistent stressors cannot be entirely excluded. HEK293 cells represent a reductionist, non-neuronal model that does not recapitulate the physiological context of HPA axis regulation, neuronal glucocorticoid receptor signaling, or FMRP expression patterns in disease-relevant cell types. While HEK293 cells endogenously express *NR3C1* and respond to glucocorticoid stimulation, validation in patient-derived neuronal models (e.g., iPSC-derived neurons from PM/FM carriers) is necessary to establish the neurobiological relevance and determine whether the observed effects generalize to the cell types most affected in FXS and FXTAS.

## Conclusions

*FMR1* CGG-repeat expansions are associated with elevated cortisol concentrations, reduced *NR3C1* expression, increased *FKBP5* expression, *NR3C1* promoter hypermethylation, and impaired glucocorticoid responsiveness in cellular models. Collectively, these findings support a model in which altered glucocorticoid signaling represents a previously unappreciated consequence of *FMR1* expansion. Although based on a limited number of molecularly confirmed carriers, and therefore requiring replication in larger independent cohorts, these observations provide a biologically plausible framework for investigating how *FMR1* CGG-repeat expansion perturbs glucocorticoid-signaling networks and highlights neuroendocrine regulation as a potential avenue for future mechanistic and therapeutic studies.

## Materials and Methods

### Cohort ascertainment

Participant recruitment was under the supervision of psychiatrists and clinical psychologists at the Institute of Mental Health and Neurosciences (IMHANS) and Shri Maharajah Hari Singh (SMHS) Hospital. The SHMS human research ethical committee granted approval, and the study was conducted in accordance with ethical standards. Participants were initially screened for FXS based on intellectual assessment and characteristic clinical features prior to molecular confirmation of the *FMR1* CGG-repeat expansion. Cognitive functioning was performed using the Wechsler Intelligence Scale for Children (WISC-V) for children aged 6– 16 years, the Wechsler Adult Intelligence Scale (WAIS-IV) for individuals ≥16 years, and the Stanford-Binet Intelligence Scales (SB-5) for younger children or those with language limitations. Full Scale IQ (FSIQ) scores were categorized using standard diagnostic criteria, and individuals with an IQ <70 were considered for further fragile X–specific testing. Comprehensive clinical evaluation included detailed prenatal and developmental history, and documentation of delays in motor, language, social, and adaptive functioning. Autism-related features were assessed using the Autism Diagnostic Observation Schedule and the Ages and Stages Questionnaire. Neurological comorbidities, including seizures and developmental regression, were recorded. Physical examination documented head circumference, dysmorphic facial features, and other phenotypic characteristics. A structured pro-forma captured demographic, clinical, and family history details.

#### Inclusion and exclusion criteria

Healthy control (CTL) participants had no history of intellectual disability, autism spectrum disorder, inherited neurological disease, or other *FMR1*-associated disorders. Individuals with acute infection, chronic inflammatory disease, autoimmune disorders, endocrine disorders known to influence glucocorticoid physiology, pregnancy, chronic corticosteroid therapy, immunosuppressive treatment, or other medical conditions likely to influence circulating cortisol concentrations were excluded whenever possible.

### Environmental stress assessment

To minimize the influence of external factors known to affect HPA-axis activity and circulating cortisol concentrations, participants were screened using a structured environmental stress questionnaire developed on previously published literature describing common psychosocial, medical, and procedural stressors that can influence glucocorticoid secretion. The questionnaire assessed recent acute stress, psychosocial stress, medication use, chronic medical conditions, and procedural stress associated with blood collection (Table 1A).

**Table 1A.** Environmental and Procedural Stress Exclusion Checklist with Supporting Literature.

| Domain | item | Yes/no | Supporting literature |
| --- | --- | --- | --- |
| Recent acute stress | Major exam or performance-related event (last 7 days) | ☐ / ☐ | Dickerson & Kemeny, 2004 |
| Psychosocial stress | Family conflict or major interpersonal stress | ☐ / ☐ | Miller et al., 2007 |
| Psychosocial stress | Caregiver-related stress | ☐ / ☐ | Lupien et al., 2009 |
| Psychosocial stress | Sleep disturbance ($\geq 3$ nights/week) | ☐ / ☐ | Meerlo et al., 2008 |
| Psychosocial stress | Change in routine or environment | ☐ / ☐ | Miller et al., 2007 |
| Medical / pharmacological | Steroid intake (past 3 months) | ☐ / ☐ | Raff, 2016 |
| Medical / pharmacological | Psychotropic medication use | ☐ / ☐ | Pariante & Lightman, 2008 |
| Medical / pharmacological | Chronic endocrine or metabolic disorder | ☐ / ☐ | Bornstein et al., 2016 |
| Procedural stress | Fear, agitation, or restraint during blood draw | ☐ / ☐ | Gunnar et al., 2009; Jansen et al., 2010 |

Each affirmative response was assigned a score of one, yielding a cumulative environmental stress score ranging from 0 to 12. Participants with a total score of 0–2 were considered eligible for inclusion, whereas those with scores >2 were excluded to minimize potential stress-related confounding. The environmental stress scores for all participants included in the study are presented in Table 1B.

**Table 1B.** Summary of environmental stress scores.

| Group (n = 10) | Score range | Median (IQR) | Participants included |
| --- | --- | --- | --- |
| Control | 0–1 | 0 (0–1) | 10/10 |
| PM | 0–2 | 1 (0–1.75) | 10/10 |
| FM | 1–2 | 1 (1–2) | 10/10 |

## Molecular Analysis

### Conventional PCR amplification of *FMR1* gene

DNA was extracted using the Tris–MgCl₂–KCl (TMK) Buffer Extraction Method from peripheral blood mononuclear cells. The samples were taken from SMHS Institute in Srinagar. PCR was used to amplify the CGG-repeat sequence within the *FMR1* gene. Two sets of repeats-specific primer pairs were used (Table 2). 4.0 units of Taq DNA polymerase (NEB), 1X Taq PCR Buffer (NEB), 0.4 mM dNTPs, 0.4μM forward and reverse primers, 1.5 mM MgCl2, and 30–60 ng of DNA were used in the 25 µl solution for the amplification. Furthermore, PCR was stabilized for GC regions by using betaine (2M). Betaine stimulates the extension reaction by binding to the major groove’s A-T pairings (Rees et al., 1993) or by binding inside the minor groove (Mytelka & Chamberlin, 1996). The samples were denaturated for 10 minutes at 98°C in a preheated thermal cycler (Bio-rad T100-thermal cycler). A final extension at 68°C for 10 minutes was then followed by 30 cycles, each lasting 40 seconds at 97°C, 35 seconds at 64°C, and 3 minutes and 40 seconds at 68°C. PCR products were analyzed by electrophoresis on 2% Ethidium Bromide-stained agarose gels.

**Table 2.**
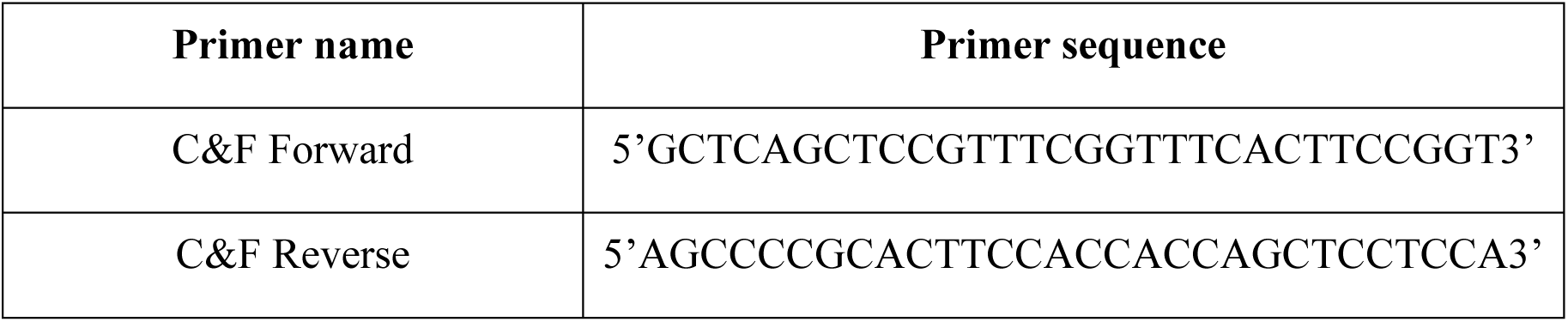
Primer sequence of C&F primers for conventional PCR.

### Bisulfite modification of DNA and methylation specific (modified) PCR

Prior to being used in methylation-specific PCR (MS-PCR), DNA was altered using a bisulfite modification kit (Zymo Research). Two primer sets were employed for this purpose (Table 3). Following Panagopoulos et al. (1999), the PCR amplification conditions were changed as follows: specifically for methylation specific PCR. A 20 μl PCR reaction was prepared with 1X buffer, 1.5 mM MgCl2, 0.2 mM dNTPs, 4U of Taq polymerase, and 0.2 μM of each primer from the sets of methylated and unmethylated primers. Following the addition of 4.0 μl of bisulfite-modified DNA to the reaction mixture, the volume was reduced to 20 μl using nuclease free water. Thermal cyclic conditions were as follow: initial denaturation at 95°C for 10 min, followed by 35cycles at 94°C for 30 sec, 66°C for 25 sec. and 72°C for 30 sec, and final extension at 72°C for 10 min. PCR products were separated by electrophoresis on 2% Ethidium Bromide-stained agarose gel.

**Table 3.**
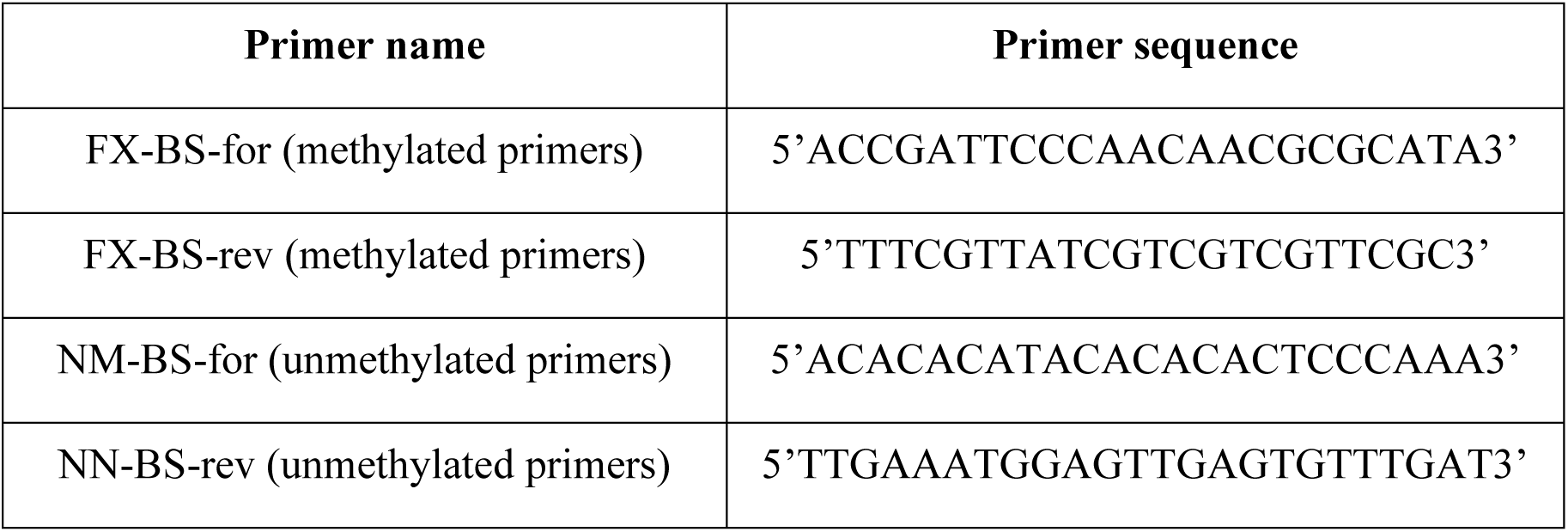
Primer sequence for methylation specific PCR.

### RNA Extraction

The TRIzol Reagent according to the manufacturer’s protocol (Invitrogen) was used in conjunction with density gradient separation and the acid guanidinium thiocyanate–phenol– chloroform technique to extract total RNA from blood-derived peripheral blood mononuclear cells (PBMCs). To isolate PBMCs by centrifugation, EDTA-anticoagulated blood was diluted with phosphate-buffered saline (PBS) and layered over Ficoll (HiMedia; HiSep^TM^ LSM1077). After collecting the interphase layer and washing it with PBS, the cell pellet was homogenized using TRIzol reagent. After centrifugation and the addition of chloroform, the RNA-containing aqueous phase was carefully moved to a new tube and precipitated using cold isopropanol. For further molecular studies, the RNA pellet was air-dried, resuspended in RNase-free water, and washed with 75% isopropanol made in DEPC-treated water.

### cDNA Synthesis

Using Revert Aid Reverse Transcriptase, which has low RNaseH activity, first-strand cDNA was synthesized from whole RNA. 500 ng of RNA was reverse transcribed in accordance with the manufacturer’s instructions in presence of random hexamers. The reaction mixture was incubated at 25°C for 5 minutes to facilitate primer annealing, followed by incubation at 42°C for 60 minutes for cDNA synthesis. The reaction was terminated by heating at 70°C for 5 minutes to inactivate the enzyme.

### Real Time PCR

Using a multicolor real-time PCR detection equipment from Bio-Rad Laboratories (CFX96^TM^ Real time system), quantitative real-time PCR (qRT-PCR) was carried out to assess mRNA expression. 2.5 µL of cDNA template was added to a final amount of 10µL for each experiment. The primer sequences used are given in Table 4.

**Table 4.**
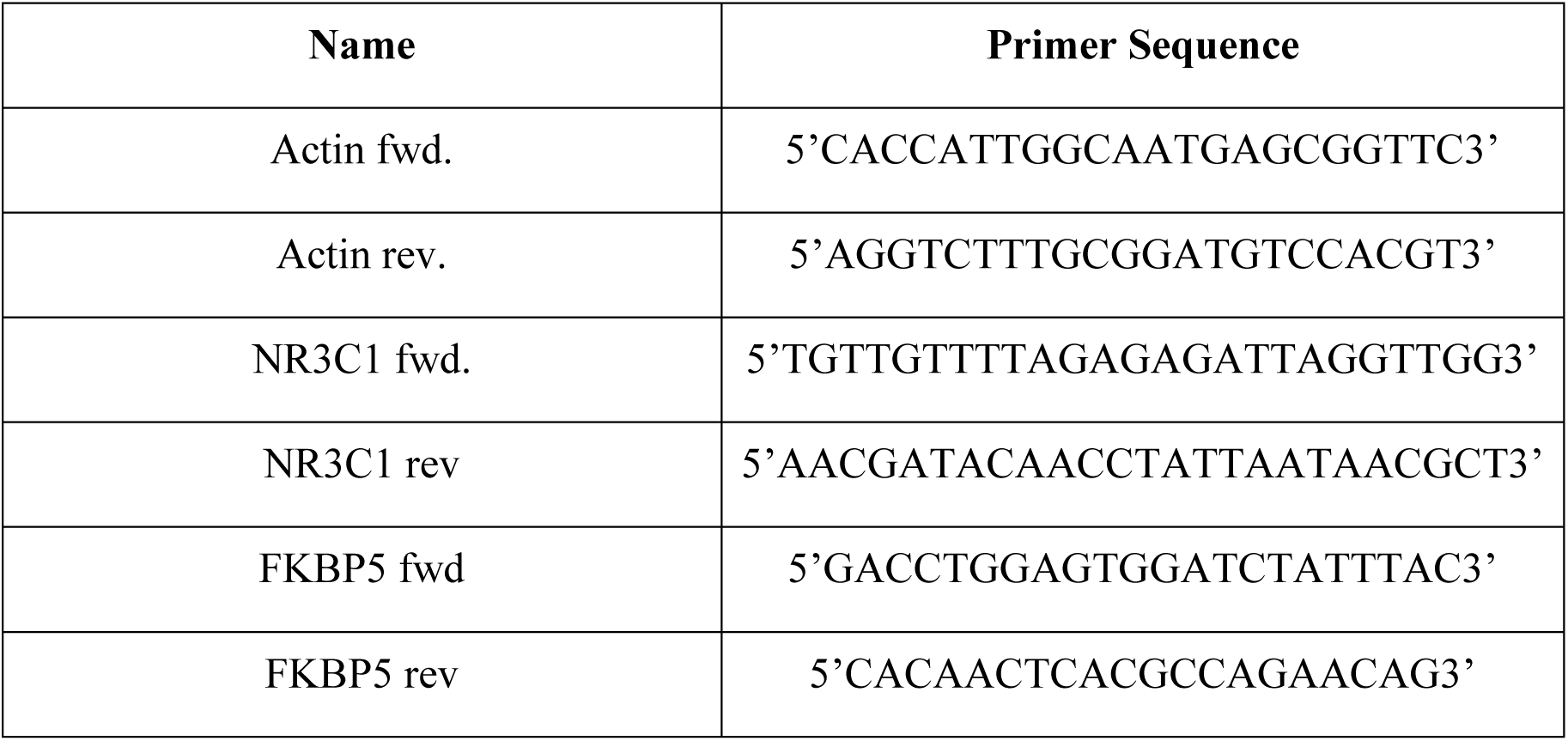
Primer sequence for Real Time qPCR.

### Enzyme Linked Immunosorbent Assay

Peripheral blood was collected in tubes containing EDTA or heparin as anticoagulants for plasma separation. Samples were centrifuged at 1000 × g for 15 minutes at 2–8°C within 30 minutes of collection. The plasma fraction was carefully aspirated and stored at −20°C or −80°C until further analysis. Repeated freeze–thaw cycles were avoided to preserve sample integrity. The ELK Human Cortisol ELISA kit was used to quantitatively estimate serum cortisol levels in accordance with the manufacturer’s instructions. Optical density (OD) was measured at 450 nm using a microplate reader. A standard curve was generated by plotting the mean OD values of the cortisol standards against their corresponding concentrations. The standard curve demonstrated a strong correlation within the assay detection range, with a coefficient of determination (R²) greater than 0.98. Cortisol concentrations in control, PM, and FM samples were determined by interpolating their OD values from the standard curve. All samples were analyzed in triplicate, and the mean OD values were used to calculate the final concentrations. Samples exhibiting a coefficient of variation (CV) greater than 15% between replicates were reanalyzed to ensure accuracy. Final cortisol concentrations were expressed in ng/mL and used for subsequent statistical analysis.

### Cell Culture, Transfection, Dexamethasone Treatment, and Gene Expression

Human embryonic kidney (HEK293) cells were maintained in Dulbecco’s Modified Eagle Medium (DMEM; Gibco, USA) supplemented with 10% fetal bovine serum (FBS) and 1% penicillin–streptomycin at 37 °C in a humidified atmosphere containing 5% CO₂. Cells were routinely subcultured upon reaching approximately 70% confluency.

For transient transfection, approximately 3 × 10⁵ cells were seeded into each well of a 6-well culture plate and allowed to attach overnight. Cells were transfected using polyethylenimine (PEI) as the transfection reagent. Expression plasmids containing either the wild-type *FMR1* and the expanded CGG-repeat construct (pcDNA3.1-CGG90) were used. An empty pcDNA3.1 vector was included as the transfection control. PEI and plasmid DNA were diluted separately in serum-free DMEM, mixed at an optimized DNA-to-PEI ratio of 1:3 (w/w), and incubated for 15–20 min at room temperature to allow complex formation. The resulting complexes were then added dropwise to the cells. Following transfection, cells were maintained under standard culture conditions. At 36 h post-transfection, cells designated for glucocorticoid stimulation were treated with dexamethasone (DEXA) at a final concentration of 1 µM. Dexamethasone was prepared as a stock solution and diluted appropriately in the culture medium to achieve the required final concentration. Cells were exposed to dexamethasone for 12 h, while corresponding untreated groups were maintained under identical conditions without dexamethasone. Thus, cells were harvested at 48 h post-transfection, corresponding to 36 h of transfection followed by 12 h of dexamethasone treatment.

For preparation of the dexamethasone stock, 25 mg dexamethasone (molecular weight 392.46 g/mol) was dissolved in 6.37 mL of appropriate solvent to obtain a 10 mM stock solution. A 200 µM working stock was subsequently prepared by diluting 20 µL of the 10 mM stock with 980 µL of solvent. For dexamethasone treatment, 10 µL of the 200 µM working stock was added to 2 mL of culture medium per well, yielding a final dexamethasone concentration of 1 µM. At the end of the treatment period, cells were harvested for RNA extraction.

### RNA extraction from HEK293 cells

Total RNA was isolated using TRIzol reagent according to the manufacturer’s instructions. Briefly, cells were lysed directly in TRIzol, followed by chloroform-mediated phase separation, isopropanol precipitation, ethanol washing, and resuspension of the RNA pellet in RNase-free water. RNA concentration and purity were determined spectrophotometrically, and equal amounts of RNA were reverse transcribed into cDNA using a commercially available reverse transcription kit. Quantitative real-time PCR (qRT-PCR) was performed to evaluate the relative expression of *NR3C1* and *FKBP5*, with actin used as the endogenous reference gene. Relative gene expression was calculated using the 2⁻^ΔΔCt^ method, with expression normalized to the untreated wild-type control group.The experimental groups included WT, vector, CGG90, WT + DEXA, vector + DEXA, and CGG90 + DEXA. All experiments were performed using at least three independent biological replicates, and data were expressed as mean ± SEM. Statistical significance was determined using one-way ANOVA followed by Tukey’s multiple-comparison test, with *p* < 0.05 considered statistically significant.

### Statistical analysis

All statistical analyses were performed using GraphPad Prism version 9.0 (GraphPad Software Inc., San Diego, CA, USA). Comparisons of cortisol concentration and gene expression among control, PM, and FM groups were performed using one-way analysis of variance (ANOVA) followed by Tukey’s multiple-comparison test. Age-stratified analyses of cortisol concentration and gene expression were carried out using two-way ANOVA, considering genotype and age group as independent variables, followed by Tukey’s post hoc test for multiple comparisons. The association between cortisol concentration and the expression of *NR3C1* and *FKBP5*, as well as the relationship between *NR3C1* and *FKBP5* expression, was evaluated using Spearman’s rank correlation analysis, and the corresponding correlation coefficient (*r*) and *p* values were documented. To assess the independent contribution of *NR3C1* and *FKBP5* expression to circulating cortisol concentrations, multiple linear regression analysis was performed using cortisol concentration as the dependent variable and gene expression levels as predictor variables.

## Acknowledgements

The authors sincerely thank all study participants and their families for their valuable participation in this study. The authors also acknowledge the Department of Biotechnology, University of Kashmir, for providing laboratory facilities, and the, Institute of Mental Health and Neurosciences (IMHANS) & Department of Psychiatry, Government Medical College Srinagar for assistance in patient recruitment and clinical evaluation. We would also like to thank DST FIST and UGC SAP-DRS-II for their support in the Department of Biotechnology, University of Kashmir. The authors would also like to thank the Qurashi laboratory members for their help and critical reading of this work.

## Author Contributions

A.Q., H.M., conceived and designed the experiments. H.M., M.M.R., A.H., performed experiments. A.Q., H.M., M.M.R., analyzed the data. A. Q, H.M., wrote the manuscript. SKH recruited the study participants and provided the blood samples.

## References.

1. Verkerk AJ, Pieretti M, Sutcliffe JS, Fu YH, Kuhl DP, Pizzuti A, et al. Identification of a gene (FMR-1) containing a CGG repeat coincident with a breakpoint cluster region exhibiting length variation in fragile X syndrome. Cell. 1991 May 31;65(5):905–14. doi:10.1016/0092-8674(91)90397-h PubMed PMID: 1710175.

2. Herman JP, McKlveen JM, Ghosal S, Kopp B, Wulsin A, Makinson R, et al. Regulation of the hypothalamic-pituitary-adrenocortical stress response. Compr Physiol. 2016 Mar 15;6(2):603–21. doi:10.1002/cphy.c150015 PubMed PMID: 27065163; PubMed Central PMCID: PMC4867107.

3. Tassone F, Hagerman RJ, Taylor AK, Gane LW, Godfrey TE, Hagerman PJ. Elevated levels of FMR1 mRNA in carrier males: a new mechanism of involvement in the fragile-X syndrome. Am J Hum Genet. 2000 Jan;66(1):6–15. doi:10.1086/302720 PubMed PMID: 10631132; PubMed Central PMCID: PMC1288349.

4. Hagerman R, Hagerman P. Advances in clinical and molecular understanding of the FMR1 premutation and fragile X-associated tremor/ataxia syndrome. Lancet Neurol. 2013 Aug;12(8):786–98. doi:10.1016/S1474-4422(13)70125-X PubMed PMID: 23867198; PubMed Central PMCID: PMC3922535.

5. Bassell GJ, Warren ST. Fragile X syndrome: loss of local mRNA regulation alters synaptic development and function. Neuron. 2008 Oct 23;60(2):201–14. doi:10.1016/j.neuron.2008.10.004 PubMed PMID: 18957214; PubMed Central PMCID: PMC3691995.

6. Sellier C, Buijsen RAM, He F, Natla S, Jung L, Tropel P, et al. Translation of Expanded CGG Repeats into FMRpolyG Is Pathogenic and May Contribute to Fragile X Tremor Ataxia Syndrome. Neuron. 2017 Jan 18;93(2):331–47. doi:10.1016/j.neuron.2016.12.016 PubMed PMID: 28065649; PubMed Central PMCID: PMC5263258.

7. Cordeiro L, Ballinger E, Hagerman R, Hessl D. Clinical assessment of DSM-IV anxiety disorders in fragile X syndrome: prevalence and characterization. J Neurodev Disord. 2011 Mar;3(1):57–67. doi:10.1007/s11689-010-9067-y PubMed PMID: 21475730; PubMed Central PMCID: PMC3057014.

8. Hall DA, Berry-Kravis E, Jacquemont S, Rice CD, Cogswell J, Zhang L, et al. Initial diagnoses given to persons with the fragile X associated tremor/ataxia syndrome (FXTAS). Neurology. 2005 Jul 26;65(2):299–301. doi:10.1212/01.wnl.0000168900.86323.9c PubMed PMID: 16043804.

9. de Kloet ER, Joëls M, Holsboer F. Stress and the brain: from adaptation to disease. Nat Rev Neurosci. 2005 Jun;6(6):463–75. doi:10.1038/nrn1683 PubMed PMID: 15891777.

10. Binder EB. The role of FKBP5, a co-chaperone of the glucocorticoid receptor in the pathogenesis and therapy of affective and anxiety disorders. Psychoneuroendocrinology. 2009 Dec;34 Suppl 1:S186–195. doi:10.1016/j.psyneuen.2009.05.021 PubMed PMID: 19560279.

11. Zannas AS, Wiechmann T, Gassen NC, Binder EB. Gene-Stress-Epigenetic Regulation of FKBP5: Clinical and Translational Implications. Neuropsychopharmacology. 2016 Jan;41(1):261–74. doi:10.1038/npp.2015.235 PubMed PMID: 26250598; PubMed Central PMCID: PMC4677131.

12. Hong J, Kapoor A, DaWalt LS, Maltman N, Kim B, Berry-Kravis EM, et al. Stress and genetics influence hair cortisol in FMR1 premutation carrier mothers of children with fragile X syndrome. Psychoneuroendocrinology. 2021 Jul;129:105266. doi:10.1016/j.psyneuen.2021.105266 PubMed PMID: 34020265; PubMed Central PMCID: PMC8217368.

13. Hartley SL, Seltzer MM, Hong J, Greenberg JS, Smith L, Almeida D, et al. Cortisol response to behavior problems in FMR1 premutation mothers of adolescents and adults with fragile X syndrome: A diathesis-stress model. Int J Behav Dev. 2012 Jan;36(1):53–61. doi:10.1177/0165025411406857 PubMed PMID: 22798702; PubMed Central PMCID: PMC3396207.

14. Seltzer MM, Barker ET, Greenberg JS, Hong J, Coe C, Almeida D. Differential sensitivity to life stress in FMR1 premutation carrier mothers of children with fragile X syndrome. Health Psychol. 2012 Sep;31(5):612–22. doi:10.1037/a0026528 PubMed PMID: 22149120; PubMed Central PMCID: PMC3434309.

15. Brouwer JR, Severijnen E, de Jong FH, Hessl D, Hagerman RJ, Oostra BA, et al. Altered hypothalamus-pituitary-adrenal gland axis regulation in the expanded CGG-repeat mouse model for fragile X-associated tremor/ataxia syndrome. Psychoneuroendocrinology. 2008 Jul;33(6):863–73. doi:10.1016/j.psyneuen.2008.03.011 PubMed PMID: 18472227; PubMed Central PMCID: PMC4408208.

16. Klengel T, Mehta D, Anacker C, Rex-Haffner M, Pruessner JC, Pariante CM, et al. Allele-specific FKBP5 DNA demethylation mediates gene-childhood trauma interactions. Nat Neurosci. 2013 Jan;16(1):33–41. doi:10.1038/nn.3275 PubMed PMID: 23201972; PubMed Central PMCID: PMC4136922.

